# GV1001 prevents ovarian follicle loss triggered by an anti-VEGF monoclonal antibody targeting non-small cell lung carcinoma xenografts in mice

**DOI:** 10.1101/719674

**Authors:** Dongmin Jang, Churl K. Min, Jisun Lee, Young-Ju Jang, Miran Kim, Kyungjoo Hwang

**Affiliations:** Department of Biomedical Sciences, Major in Molecular Medicine, Graduate School of Medicine, Ajou University, Suwon, South Korea; Department of Biological Sciences, Ajou University, Suwon, South Korea; Department of Obstetrics and Gynecology, Ajou University Medical Center, Suwon, South Korea

**Author notes:** Department of Obstetrics and Gynecology, Kyungpook National University Hospital, Daegu, South Korea (Jisun Lee).

**Keywords:** Premature ovarian failure, Combinatorial chemotherapy, GV1001, Bevacizumab, Non-small cell lung carcinoma, Tumor xenografts

## Abstract

Premature ovarian failure (POF) that could result from chemotherapy applied to young female cancer patients is a significant challenge in reproductive biology. It is widely believed that the hyperactivation of dormant primordial follicles following chemotherapy is a leading cause of POF, but it remains unclear how therapeutic cues are generated and transduced into follicular activation. Here, we provide evidence that supports that GV1001, an immunotherapeutic peptide targeting telomerase, plays a role in the deterrence of POF in mice. I*n vivo* non-small cell lung carcinoma (NSCLC) tumor xenografts were produced by inoculating NSCLC cells into the flank of BALB/c female athymic mice and then subjected to cancer chemotherapy with GV1001 and bevacizumab, an anti-cancer antibody drug, humanized anti-VEGF monoclonal antibody. Bevacizumab when administered at the dosage of 5 mg/kg for three weeks was effective in inhibiting growth of NSCLC tumor xenografts, and its anti-cancer efficacy was not interfered by the presence of GV1001. As expected, bevacizumab induced follicular loss by accelerating primordial follicle growth into primary or secondary follicles concomitant with a decline of serum antimullerian hormone (AMH) level and deactivation of Foxo3 signaling as evidenced by immunohistochemical and immunofluorescent assessment. However, bevacizumab-induced follicle stimulating effects were mitigated by GV1001 co-administration as evidenced by the analysis of follicular count, serum AMH level, and Foxo3a expression. From this study, we propose that a combinatorial administration of GV1001 and bevacizumab could deter POF of young female cancer patients without hampering the anti-cancer effectiveness of bevacizumab.

## INTRODUCTION

Premature ovarian failure (POF) is one of the most serious complications of chemotherapy administered to young female cancer patients. The toxicity of chemotherapeutics depends on chemotherapy regimen, dosage and patient age. Radiation applied to the pelvic area along with alkylating agents, e.g. cyclophosphamides, administered to patients are known to have the most detrimental effects on ovarian reserves [4]. In order to overcome any ovo-toxic effect due to radiation therapy or chemotherapy, cryopreservation of oocytes, embryos or ovarian tissues are strongly suggested as available options for fertility preservation. Given that fertility preservation methods require a series of endeavors in a precise time frame-dependent manner, including ovulation induction and partner participation, they may not be suitable for all cancer patients. Rather, as a replacement for invasive cryopreservation techniques, some studies have now focused on finding a new agent that can be co-applied at the time of chemotherapy aimed at a potential decrease in ovo-toxicity of chemotherapy agents. Some agents have been studied to prevent chemotherapy-induced ovarian damage. Imatinib, a competitive tyrosine-kinase inhibitor, and sphingosine-1-phosphate (S1P), an inhibitor of the ceramide-promoted apoptotic pathway, for example, have been ascribed to anti-apoptotic activity, and AS101 is known to reduces apoptosis in granulosa cells in growing follicles thereby preventing cyclophosphamide-induced primordial follicle loss [5].

GV1001 was first developed as an anti-cancer peptide vaccine derived from human telomere reverse transcriptase, targeting telomerase [6], based on the fact that telomerase is expressed robustly in most cancer cells but found scarcely in normal cells. Several phase I/II trials have been conducted on patients of pancreatic cancer, advanced melanoma, or non-small cell lung cancer and met with some promising anticancer effects [3, 7]. When used along with other chemotherapeutic agents, GV1001 was able to induce T cell-mediated anti-cancer effects. GV1001 was also evaluated to have protective effect against renal ischemia injury by reducing inflammation and apoptosis [2].

Despite multiple extra-telomeric actions of GV1001, most studies have focused on anticancer effects of GV1001 thus lacking systematic studies on any restorative effect of GV1001 against chemotherapy-induced ovary dysfunction. Another targeted therapeutic agent bevacizumab (Avastin), a humanized anti-VEGF monoclonal antibody [9], has been widely used in combination with other chemotherapeutics against many cancers including metastatic colorectal cancer, non-small cell lung cancer, breast cancer, and ovarian cancer [1, 8]. The objective of our study, therefore, was to investigate whether a combinatorial administration of GV1001 and bevacizumab has any beneficiary effect on ovarian functions, particularly, on the loss of ovarian follicles without hampering their anticancer activities, and thereby to provide a possible combinatorial chemotherapy regimen with broad relevance for POF control.

## MATERIALS AND METHODS

### Animals

Five-week-old female BALB/c-nu mice (n = 80) were purchased from Oriental Bio (Seongnam, S. Korea) and allowed to acclimatize to and recover from moving-related stress for at least 7 days prior to the study. Mice were allowed free access to chlorinated water and irradiated foods and kept in climate-controlled 12 hr light/dark cycle with daily visual monitoring. All animal experiments were pre-reviewed and approved by the Institutional Animal Research Ethics Committee at Ajou University (https://eirb.ajoumc.or.kr) and conducted in the Laboratory Animal Research Center at Ajou University Hospital, Suwon, S. Korea (http://larc.ajoumc.or.kr) under the supervision of a certified veterinarian (IACUC No. 2016-0023).

### Cell lines and cell cultures

Human non-small cell lung cancer (NSCLC) cell lines H441 were purchased from American Type Culture Collection (#HTB-174, Manassas, VA) and maintained in RPMI - 1640 media supplemented with 2 mM L-glutamine, 10 mM HEPES, 1 mM sodium pyruvate, 4.5 g/L glucose, 1.5 g/L sodium bicarbonate, and 10% FBS (Gibco-Invitrogen, Grand Island, NY) at 37°C under 5% CO_2_.

### Tumor xenografts in athymic mice and GV1001 treatment

About 200 μl of NSCLC cells suspension (3~5 × 10^6^ cells/ml) was injected subcutaneously (*s.c*.) into the left flank of an athymic mouse. Tumor-bearing mice were randomly assigned to different treatment (n=5 for each treatment) when the mean tumor volume reached 100~200 *mm*^3^in size. Then, the mice were administered s.c. either with bevacizumab (Roche Pharma, Basel, Switzerland) at 5 mg/kg, with GV1001 (GemVax & KAEL, Seongnam, S. Korea) at 0.5 or 2 mg/kg, or with both bevacizumab at 5 mg/kg and GV1001 at 2 mg/kg twice a week for 3 weeks while control mice received equal volume of PBS. The injection volume was 50 *μℓ*. Tumor diameters were measured externally with a vernier caliper once a week, and tumor volume (V) was calculated by the equation: 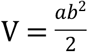 where *a* = length and *b* = width [10].

### Ovarian histology and follicle counting

The mice were sacrificed at the end of the experiment by cervical dislocation, and the ovaries were excised surgically at necropsy. The excised ovaries were fixed, dehydrated, and embedded in paraffin using an automated tissue processor (Excelsior, ThermoFisher Sci., Waltham, MA) according to a standard protocol. Sections of 5 *μ*m-thick were cut with a microtome before mounted on Superfrost-Plus slides (ThermoFIsher Sci.). Haematoxylin and eosin staining was performed using a Leica Autostainer XL (Leica Biosystems, Wetzlar, Germany). All the subsequent tissue processing was performed at the Pathology Department of Ajou University Hospital. Briefly, tissue sections were deparaffinized, rehydrated, and mordanted in Bouin’s solution for 15 min at 60℃. Then, slides were washed for 5 min in running tap water prior to being placed in Weigart’s haematoxylin for 10 min. Slides were again washed for 5 min in running water, followed by staining in Biebrich Scarlet-Acid Fuchsin solution for 5 min. After brief rinsing in H_2_O, slides were transferred to phosphotungstic/phosphomolybdic acid for 10 min before stained in aniline blue for 5 min. Slides were briefly rinsed in H_2_O, fixed in 1% acetic acid for 1 min, dehydrated, cleared, and dried before mounted on a microscope for viewing. Early follicular developmental stage was histologically classified according to previously accepted criteria [11, 14] as follow: A primordial follicle was defined as an oocyte surrounded by a single layer of flattened squamous pregranulosa cells. A primary follicle was defined as an oocyte surrounded by a single layer of cuboidal granulosa cells. Occasionally, follicles appeared as an intermediate stage between the primordial and primary stages, with both cuboidal and squamous granulosa cells. If the cuboidal cells predominated, the follicle was classified as a primary follicle. A secondary follicle has two or more layers of cuboidal granulosa cells with no visible antrum. An antral follicle is defined as a follicle with a fluid-filled antral space. A preovulatory follicle is the largest follicle with a cumulous granulosa cell layer. For statistical analysis, the antral follicle and the preovulatory follicle were grouped together as antral follicles as suggested by Chang *et al*. [15].

### Immunohistochemistry

The excised ovaries were paraformaldehyde fixed, paraffin embedded, and subjected to microtomy according to standard procedures as described in *Histological follicle assessment*. About 5 *μ*m-thick paraffin sections on glass slides were deparaffinized, then rehydrated before quenching the endogenous peroxidase activity by incubation in 3% hydrogen peroxide in methanol for 30 min. Slides were then subjected to antigen retrieval by boiling in 10 mM citrate buffer, pH 6.0, for 20 min and subsequent blocking with 10% nonspecific immune serum in the blocking solution for 1 hr before incubation overnight at 4℃ with anti-Foxo3a antibodies (Santa Cruz Biotech, Santa Cruz, CA) at 1:100 dilution in the antibody diluent. DAB color reaction was performed with the Histostain-Plus Bulk Kit (Invitrogen, Carlsbad, CA) followed by DAPI counterstaining before microscopic examination under an Olympus microscope D50 (Tokyo, Japan).

### Immunofluorescence

The ovaries were fixed, paraffin-embedded, and thin-sectioned as described in *Immunohistochemistry* before subjected to immunofluorescence staining according to a standard protocol using following antibodies: primarily with anti-Foxo3a antibodies (Santa Cruz Biotech.) diluted at 1:100 and secondarily with Alexa 488-congugated goat anti-rabbit Ig (Invitrogen) diluted at 1:400. DAPI counter-staining was used to control the process of immunofluorescence. After staining, the tissue sections were observed with LSM 510 META confocal laser-scanning microscope (Carl Zeiss, Oberkochen, Germany).

### ELISA assay

The concentration of serum antimullerian hormone (AMH) was determined using the ELISA assay kit (Anshlabs, Webster, TX) according to the provided description with minor modification. In brief, undiluted serum samples were dispensed into the anti-AMH antibody coated 96 well plate, and anti-AMH detection antibodies labeled with biotin was added. After washing of samples, 100 *μℓ* of streptavidin-enzyme conjugate-RTU was added, followed by 100 *μℓ* of substrate solution containing 3,30,5,50-tetramethylbenzidine (TMB) and incubation for 8-12 min at room temperature. A microplate spectrophotometer (model SpectraMax 190, Molecular Devices, Sunnyvale, CA) was used to measure the absorbance at 450 nm and 630 nm. The absorbance was converted into the enzymatic turnover of substrates from the calibration curve provided.

### Statistical analysis

All experiments were repeated at least three times. Quantitative variables are given as means ± standard deviations. Statistical significance of the data was carried out with the *Student’s* t-test with P values at <0.05.

## RESULTS

### Bevacizumab induces follicle loss by accelerating dormant follicle activation

First, we ascertain whether bevacizumab causes follicular loss in mice. Fig 1A illustrates representative views of the formalin-fixed, paraffin-embedded, and eosin/haematoxylin stained mouse ovary section before and after the chemotherapy. Bevacizumab at 5 mg/kg drastically reduced the number of primordial follicles, which are indicated by asterisks, in the mouse ovary. Contrarily, the reduced number was smaller in the primordial follicles treated with GV1001 at either 0.5 mg/kg or 2 mg/kg.

**Fig 1.**
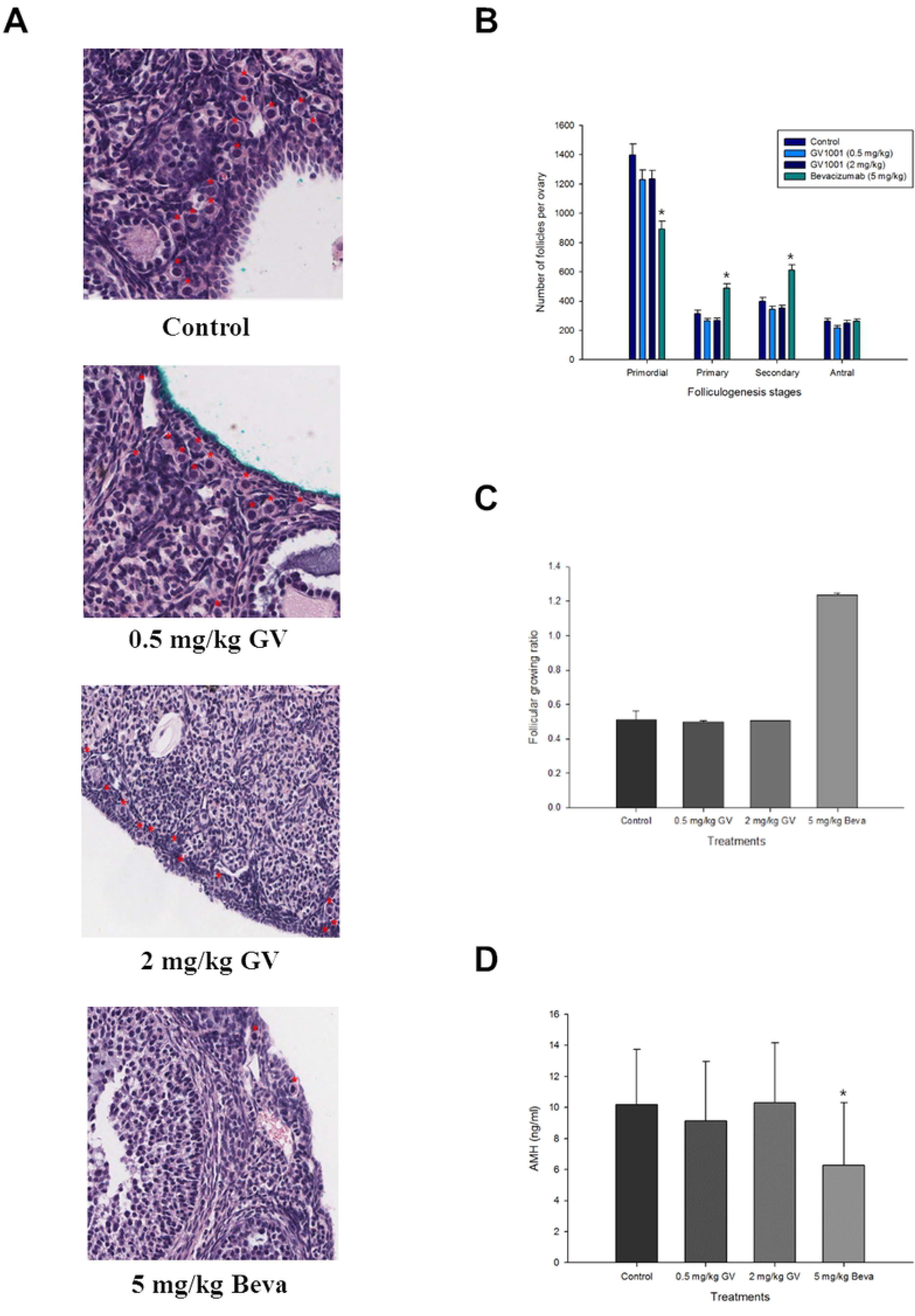
Histological evaluation of ovarian follicles after exposure to bevacizumab and/or GV1001. (A) Representative histological ovary tissue section images after haematoxylin and eosin staining. NSCLC tumor-bearing mice were s.c. administered with PBS (controls), with GV1001 at 0.5 mg/kg, with GV1001 at 2 mg/kg, or with bevacizumab at 5 mg/kg for 3 weeks before the ovaries were excised, fixed, paraffin-embedded, micro-sectioned, and subjected to haematoxylin and eosin staining. The asterisks denote the primordial follicles. GV, GV1001; Beva, bevacizumab. (B) Ovarian follicles at each early developmental stage (primordial, primary, secondary and antral follicles) were counted under the microscope according to criteria as described in *Materials and Methods*. *p<0.05 against controls at each folliculogenesis stage. (C) Follicular growing ratio was calculated according to the equation described in *Materials and Methods*. GV, GV1001; Beva, bevacizumab. (D) The serum level of antimullerian hormone was measured using commercial ELISA kits as described in *Materials and Methods*. GV, GV1001; Beva, bevacizumab. *p<0.05 against controls.

A more stringent test of toxicity on the ovarian follicles would be to assess a change in the number of follicles at each development stage after exposure to chemotherapy. We therefore counted the number of follicles at each developmental stage (primordial, primary, secondary, and antral) in ovary sections under the microscope. The results clearly showed a trend towards a reduction in the number of primordial follicles when bevacizumab or GV1001 was administered compared to controls. However, the magnitude of reduction was significantly bigger in the presence of Bevacizumab at 5 mg/kg compared to GV1001 at either 0.5 mg/kg or 2 mg/kg. There were little difference in GV1001, 0.5 mg/kg *vs* 2 mg/kg, in terms of primordial follicular number. On the contrary, there seemed to be no apparent decrease in the number of primary, secondary, or antral follicles under any chemotherapy regimen compared to each control. Rather, the numbers of primary and secondary follicles were increased by bevacizumab treatment (Fig 1B). This observation could be interpretated as that whereas the number of total follicles per ovary were decreased by the chemotherapy, the follicles at different developmental stages was not proportionally affected. The antral follicle was affected to a minimum by the chemotherapy regimens used in this study while the numbers of primary and secondary follicles were rather increased by bevacizumab treatment. To emphasize the developmental progression of the primordial follicles to the primary or the secondary follicles, we sought out follicular growing ratio, which was obtained from 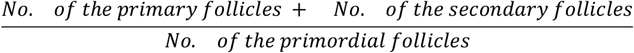. Compared to either control or GV1001 treatment, bevacizumab treatment exhibited about 2.5-fold increase in follicular growing ratio (Fig 1C). This result implies that unlike GV1001, bevacizumab may trigger follicular activation from the dormant primordial follicles, which could result in the increased follicular growing ratio by bevacizumab treatment. AMH is a negative regulator of follicle activation, produced by granulosa cells of small growing follicles [21]. As evident in Fig 1D, the AMH concentration in the sera of mice after exposure to bevacizumab was about 40% lower than control or the mice exposed to 2 mg/kg of GV1001, confirming that the released AMH-mediated suppression of the primordial follicles could be implicated in primordial follicle loss as suggested previously [16].

### Co-administration of GV1001 deters follicle activation

Next, we compared the extent of follicle activation resulting from co-administration of GV1001 and bevacizumab to that from single administration of bevacizumab to address if there was any protective effect of GV1001 on follicle activation/growth and thus follicle loss. When GV1001 was co-administered with bevacizumab, there was a significant reduction in the loss of primordial follicle induced by bevacizumab. Fig 2A illustrated that the bevacizumab-induced decrease in the number of primordial follicles was recovered almost to the control level upon co-administration with GV1001 at either 0.5 mg/kg or 2 mg/kg. The rescue action of GV1001 was also evident on primary and secondary follicles. The bevacizumab-triggered increase in the number of primary and secondary follicles also returned to the control level upon GV1001 co-administration. One generalization permitted from the present evidence is that GV1001 might deter the growth of primordial follicles to primary and/or secondary follicles triggered in the presence of bevacizumab. The magnitude of follicle activation, which is accounted for by follicular growing ratio, was reduced by as much as 58% when GV1001 was co-administered with bevacizumab relative to bevacizumab alone (Fig 2B). More consistently, the lower serum AMP level induced by the presence of bevacizumab also returned to the control level when GV1001 was co-administered (Fig 2C), confirming the involvement of AMH-mediated suppression in the bevacizumab-induced follicle activation.

**Fig 2.**
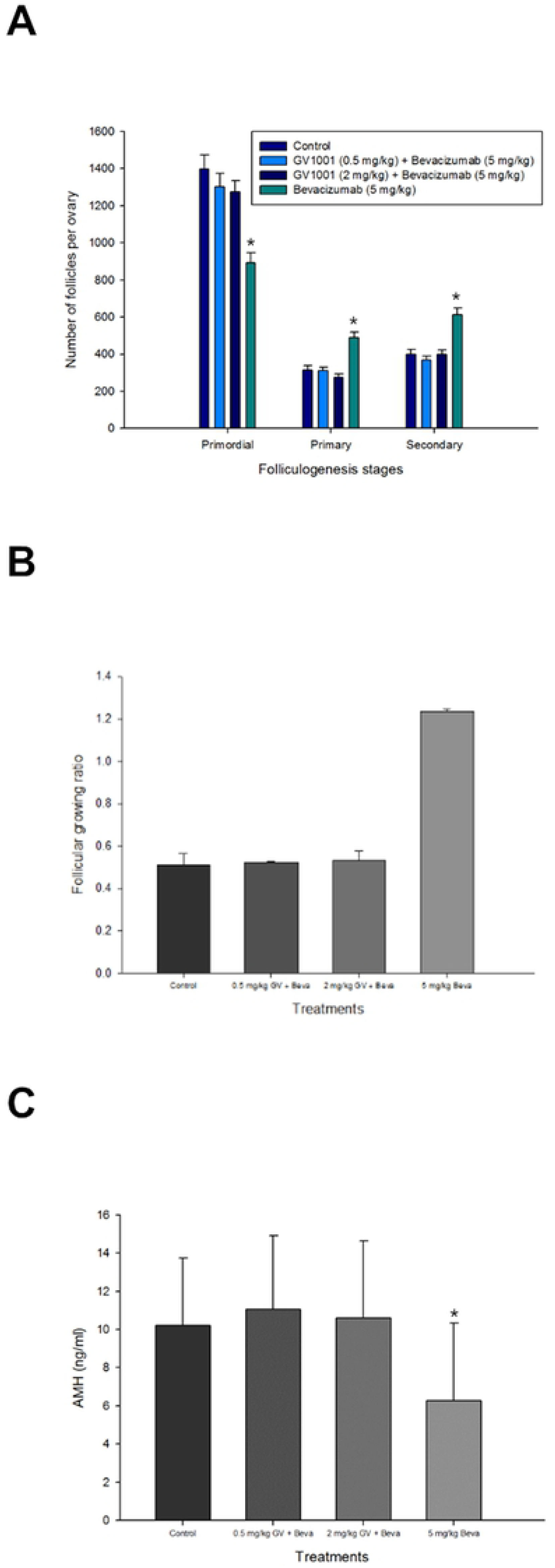
Deterrence of bevacizumab-induced ovarian follicle loss by GV1001. (A) NSCLC tumor-bearing mice were s.c. administered with PBS (controls), with GV1001 at 0.5 mm/kg plus bevacizumab at 5 mg/kg, with GV1001 at 2 mg/kg plus bevacizumab at 5 mg/kg, or with bevacizumab at 5 mg/kg for 3 weeks before the ovaries were excised, fixed, paraffin-embedded, micro-sectioned, and subjected to haematoxylin and eosin staining. The number of each growing ovarian follicles were counted according to the criteria described in *Materials and Methods*. *p<0.05 against bevacizumab at each folliculogenesis stage. (B) The follicle growing ratios were calculated based on the data in (A). (C) The serum level of antimullerian hormone was measured as described in Figure 1. *p<0.05 against bevacizumab.

### Foxo3 signaling is involved in bevacizumab-induced follicular activation

The forkhead transcription factor Foxo3 is the transcription factor that has been implicated in diverse biological processes, including metabolism, cellular stress responses, and aging. More to the point, Foxo3 serves an essential role in the regulation of follicle growth as a suppressor of follicular activation [13, 16]. In this regard, it is expected that the expression level of Foxo3 would be decreased to predispose follicles to mature in the presence of bevacizumab and rescued to the normal level in the co-presence of GV1001. To understand how Foxo3 regulates ovarian follicle growth, both immunohistochemical and immunofluorescent localization of Foxo3a in primordial follicles were observed under different chemotherapy regimens. Foxo3a expressions in primordial follicles were lower in bevacizumab-received mice than either in control mice or in GV1001-received mice (Fig 3A). In support of this notion, the bevacizumab-induced reduction in Foxo3a expression was likely to be rescued upon GV1001 co-administration in immunofluorescence analysis (Fig 3B).

**Fig 3.**
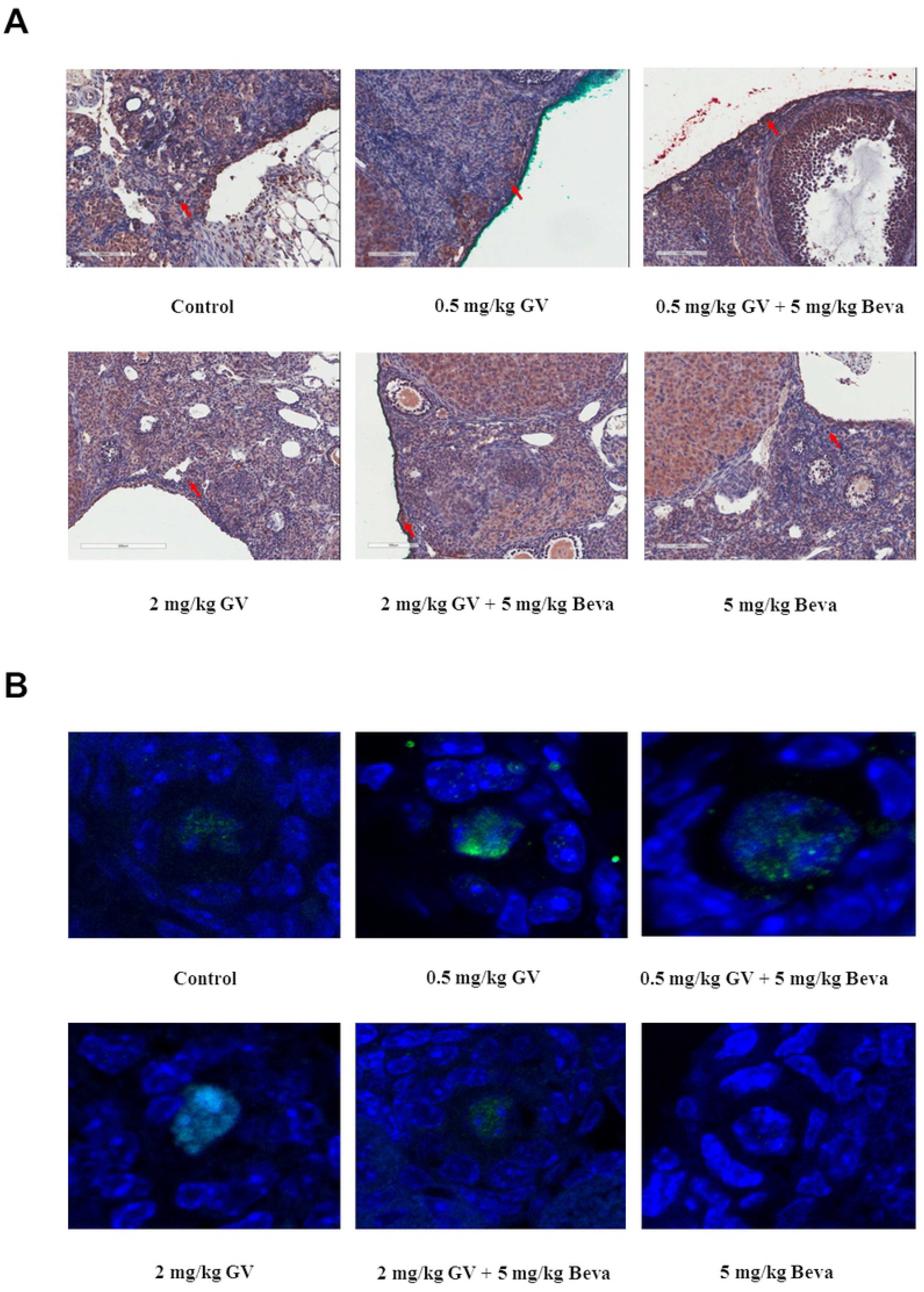
Expression and localization of Foxo3a in ovarian primordial follicles. (A) Immunohistochemical localization of Foxo3a in the ovarian primordial follicles under different chemotherapy regimens. The ovaries of NSCLC tumor-bearing mice after chemotherapy was excised, fixed, paraffin-embedded, and thin-sectioned to visualize the ovarian follicles and subjected to immunostaining with anti-Foxo3a antibodies. DAPI was used for the nuclear counter-staining. Arrows indicates the primordial follicles. GV, GV1001; Beva, bevacizumab. (B) Immunofluorescence analysis of Foxo3a expression in ovarian primordial follicles. The ovaries were dissected and processed as in (A) and subjected to immunofluorescent staining as described in *Materials and Methods*. Green and blue fluorescence indicates Foxo3a and the nuclei, respectively. GV, GV1001; Beva, bevacizumab.

### GV1001 sensitizes non-small cell lung cancer cells in xenografts using athymic nude mice to bevacizumab immunotherapy

To determine whether GV1001 interferes with the anticancer effects of bevacizumab, we tested its activity *in vivo* on a cancer xenograft model. Bevacizumab is commonly used in immunotherapy protocols for lung cancer. Thus, for this study, the cell line used for cancer xenograft model is non-small cell lung cancer (NSCLC). By injecting NSCLC cells into the flank of athymic nude mice, we produced NSCLC tumor xenografts. Then, the tumor-bearing mice were administered *s.c.* with bevacizumab or in combination with GV1001 to detect any change in tumor xenograft growth. The tumor-bearing mice under present chemotherapy regimens appeared outwardly normal and did not show abnormal weight gain or statistically significant differences in food intake and behavior (data not shown), suggesting little tumor burden or toxicity due to the chemotherapy. The volume increase between the tumor volumes before and after the PBS injection as control were 973.3 ± 16.1 mm^3^ (n=5). On the other hand, bevacizumab administered alone (5 mg/kg) caused a significant decrease in the tumor volume increase to 264.7 ± 1.9 mm^3^ (n=5) compared with control. Also, bevacizumab co-administered with GV1001 (2 mg/kg) caused a significant decrease in tumor volume increase to 179.9 ± 1.0 mm^3^ (n=5) compared with control. The tumor growth inhibition rate, which was defined as the percentage of tumor growth inhibition relative to controls, was 72.80% and 81.50% in bevacizumab single administration and in GV1001/bevacizumab co-administration, respectively (Table 1). Our results show that there is a potential of synergic effect about anti-cancer therapy in GV1001/bevacizumab co-administration. We also think this synergic effect appears regardless of the original function of GV1001, which is known as anti-cancer peptide vaccine by activating the T cell immune system, because mice used in the experiment are the athymic mice having a greatly reduced number of T cells.

**Table 1.** GV1001 increases sensitivity of Non-small cell lung carcinoma (NSCLC) to bevacizumab immunotherapy in vivo.

|  | No. of mice examined (n) | Initial tumor volume (mm<sup>3</sup>) | Final tumor volume (mm<sup>3</sup>) | Volume increase (mm<sup>3</sup>) | Inhibition rate (%) |
| --- | --- | --- | --- | --- | --- |
| Controls | 5 | 261.4 ± 0.3 | 1234.7 ± 15.9 | 973.3 ± 16.1 | - |
| Bevacizumab | 5 | 226.6 ± 1.2 | 491.3 ± 0.7 | 264.7 ± 1.9 | 72.8 |
| Bevacizumab + GV1001 | 5 | 385.3 ± 0.8 | 565.3 ± 0.2 | 179.9 ± 1.0 | 81.5 |

After the tumor reached visible size, mice were subjected to the s.c. drug injection as follows: controls (PBS), bevacizumab (5 mg/kg), and bevacizumab (5 mg/kg) and GV1001 (2 mg/kg) co-administration. The initial and final tumor volume indicate the tumor volume measured before and after drug administration.

## DISCUSSION

These studies demonstrate that the bevacizumab-induced follicle loss in mice are mainly due to hyperactivation or awakening of primordial follicles from dormancy to initiate early follicular growth, progressing to advanced stage follicles concomitant with a decline of primordial follicles. In accordance, we were able to detect concurrent abnormalities in both AMH secretion and Foxo3a expression in primordial follicles, both of which are known to act as suppressors of ovarian follicle activation. These findings further support the importance of Foxo3 in the regulation of chemotherapy-induced POF, raising intriguing possibility that manipulation of Foxo3 could be a novel means for the deterrence of POF in human.

The functional life span of the ovaries is determined mainly by the number of oocytes present, a number that is known to decline precipitously during both fetal development and postnatal life. Furthermore, it is well documented that exposure of the ovaries to a variety of pathological insults, such as anticancer therapies and environmental toxicants, accelerates oocyte and follicle depletion, consequently hastening the time at which ovarian failure is observed. Accordingly, over the past several years a tremendous amount of research effort has been expended to uncover the genetic and molecular mechanisms responsible for determining the size of the follicle reserve endowed at birth as well as the rate at which this stockpile of follicles is subsequently depleted [17, 18].

In this study, we employed a histological evaluation of the ovary to classify developmental stage-specific follicles since this methodology has been the most accurate, albeit tedious, means to provide an estimate of the number of follicles in the ovaries, in particular, follicles at early stages of growth (primordial, primary, secondary and antral) [11, 14]. A primordial follicle was defined as an oocyte surrounded by a single layer of flattened squamous pre-granulosa cells. A primary follicle was defined as an oocyte surrounded by a single layer of cuboidal granulosa cells. Occasionally, follicles appeared as an intermediate stage between the primordial and primary stages, with both cuboidal and squamous granulosa cells. If the cuboidal cells predominated, the follicle was classified as a primary follicle. Secondary follicles had two or more layers of cuboidal granulosa cells with no visible antrum. Antral follicles were defined as follicles with a fluid-filled antral space.

Recent view regarding chemotherapy-induced follicular loss suggests that many chemotherapeutic agents trigger an initiation of dormant follicle growth, occurring concurrently with large follicle apoptosis [19]. Destruction of large follicles causes a reduction of AMH secretion and thus a loss of suppression of the primordial follicle reserve. The loss of suppression would then result in the activation of primordial follicles to replace the dying cohort of growing follicles [20]. This line of speculation prompted us to examine whether either GV1001 or bevacizumab chemotherapy stimulated the early progression of ovarian follicle growth, causing concurrently the numbers of follicles at more advanced stages to increase while a corresponding decrease in primordial follicles.

Primordial follicles are long-lived structures assembled early in life. The mechanisms that control the balance between the conservation and activation of primordial follicles are critically important for fertility and dictate the onset of menopause [13]. In this study, we showed that mice that were exposed to bevacizumab chemotherapy exhibited a distinctive ovarian phenotype of global follicular activation leading to oocyte death and early depletion of functional ovarian follicles. At the same time, bevacizumab caused a decline in AMH serum level and the deactivation of Foxo3a in primordial follicles. These results signify the possibility that accelerated follicular initiation plays a major role in premature ovarian failure by depletion of primordial follicle and consequent ovarian failure, a common cause of infertility and premature aging in women. Another consideration implied by these findings is that there are repressive feedback mechanisms that restrict early follicle growth, and this restriction could be damaged by anti-cancer chemotherapy as reflected by abnormalities in serum AMH levels and Foxo3 signaling.

Of particular interest is the fact that an exceptional phenotype of athymic mice associated with co-administrated medications (GV1001 and bevacizumab), which therefore we took advantage of, presents a unique opportunity to assess functional consequences of immunotherapy differentially: one that is associated with anticancer activity from the other associated with ovarian follicle activation. GV1001 was developed as an immunotherapeutic peptide targeting telomerase, thus enabling to induce T cell mediated anti-cancer effects. However, our experiment cannot be expected T cell mediated anti-cancer effects, because our experiment was used to athymic nude mice characterized by absence of the thymus. Thus, we suggest another mechanism increasing anti-cancer effect through GV1001/bevacizumab co-administration, such as apoptosis by anti-angiogenesis [25].

Athymic mice are characterized by deteriorated or absence of the thymus and thus by nonfunctional T cells, which enables precise perturbation of T cell mediated anticancer activities of GV1001 while leaving its implication with follicular activation unaffected. Our data clearly demonstrate that GV1001 does not interfere with or even sensitize the bevacizumab-mediated anticancer activity against NSCLC carcinoma in athymic mice when co-administrated.

More to the point, Foxo3a deactivation as indicated by its reduced histological/cytological expression in primordial follicles after exposure to bevacizumab appear to be in line with the view that PTEN/PI3K/AKT/FOXO3 signaling pathway could be harnessed in the regulation of follicular activation/growth as previously suggested [16]. Further supporting evidence includes (*i*) PTEN-specific mutation in primordial follicles results in excessive activation of the entire pool of primordial follicles, which in turn leads to premature depletion of all primordial follicles [22]; (ii) FOXO3a -/- mice reveal up-regulation of primordial follicle activation [23]. One interesting question that remains to be answered would be the functional correlation between FOXO3 signaling and vascular endothelial growth factor A (VEGF-A)-derived signaling. Recently, it has been well documented that there exist multiple isoforms of VEGF-A resulting from alternative splicing, and depending upon the amino acid sequences, some act as proangiogenic factor while others act as antiangiogenic factor. Proangiogenic VEGF-A isoforms appear to promote initial recruitment and early development of ovarian follicles whereas antiangiogenic VEGF-A isoforms appear to suppress these processes [24]. Obviously, bevacizumab derived from anti-VEGF antibodies does not differentiate VEGF isoforms, but it is a quite intriguing speculation that the balance between proangiogenic isoforms and antiangiogenic isoforms might be disturbed by bevacizumab administration and thus influences early follicle development as suggested by McFee et al. [24]. The PTEN/PI3K/AKT/FOXO3 signaling pathway is, therefore, highly likely to be richly endowed with the versatilities needed to confer on cells the ability to cope with diverse internal or external stresses, including one from chemotherapy

Given the paucity of data on this issue, more studies are needed to fully delineate the underlying mechanisms of the chemotherapy-associated primordial follicle activation. At this moment, it suffices to say that it is tempting to speculate that the present findings may be of broad relevance for cancer chemotherapy-related POF in the future. Nevertheless, it has been shown that GV1001 is effective in preserving ovarian function when co-administered with an anticancer agent. Taken together, a potential therapeutic regimen consisting of GV1001 and bevacizumab co-administration seems to be a reliable means by which one can deter the progression of cancers while preserving ovarian follicle reserve in young female cancer patients.

## AUTHOR CONTRIBUTIONS

Conceived and designed the experiments: DMJ MRK. Performed the experiments: DMJ JSL. Analyzed the data: DMJ MRK. Contributed reagents/materials/analysis tools: DMJ CKM YJJ MRK KJH. Wrote the paper: DMJ CKM YJJ.

